# Anti-apoptotic and anti-inflammatory properties of grapefruit IntegroPectin on human microglial HMC3 cell line

**DOI:** 10.1101/2023.07.20.549931

**Authors:** Miriana Scordino, Giulia Urone, Monica Frinchi, Chiara Valenza, Angela Bonura, Chiara Cipollina, Rosaria Ciriminna, Francesco Meneguzzo, Mario Pagliaro, Giuseppa Mudò, Valentina Di Liberto

**Author notes:** **Corresponding Author Valentina Di Liberto** - Dipartimento di Biomedicina, Neuroscienze e Diagnostica Avanzata, Università di Palermo, corso Tukory 129, 90134 Palermo, Italy. These authors contributed equally to this work (Miriana Scordino, Giulia Urone). These authors contributed equally to this work (Giuseppa Mudò, Valentina Di Liberto).

## Abstract

**Background and aim:** 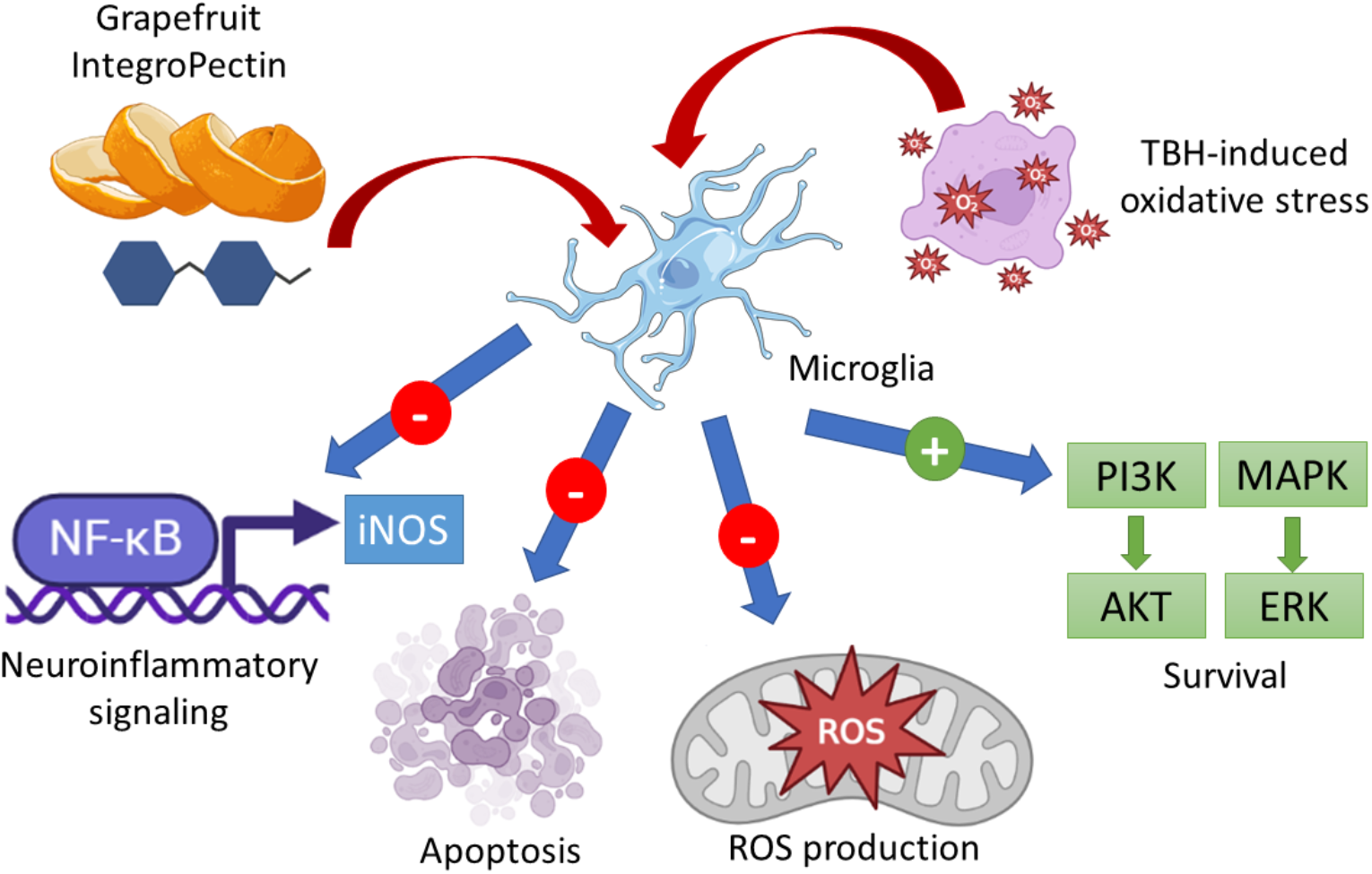

Despite the intense research, most therapeutic strategies failed in preventing or treating neurodegenerative diseases, characterized by a combination of chronic neurodegeneration, oxidative stress and neuroinflammation. The broad protective activity of IntegroPectin derived from industrial waste grapefruit peel via hydrodynamic cavitation has been recently characterized. In this study, we investigated the beneficial effects of grapefruit IntegroPectin treatment in microglia cells exposed to oxidative stress conditions.

**Experimental procedure:** Human microglial HMC3 cells were challenged with tert-butyl hydroperoxide (TBH), a powerful hydroperoxide, in the presence of grapefruit IntegroPectin. The apoptotic process, the oxidative stress and the neuroinflammatory responses with the relative intracellular cascades were evaluated.

**Key results:** Grapefruit IntegroPectin fully counteracted the apoptotic process induced by cell exposure to TBH. The protective effects of grapefruit IntegroPectin were accompanied with a decrease in the amount of ROS, and were strictly dependent on the activation of the PI3K/Akt cascade. Finally, IntegroPectin treatment inhibited basal microglia activation and the neuroinflammatory response by down-regulating the PI3K-NF-kB-iNOS cascade.

**Conclusions and implications:** These findings reveal that the innovative IntegroPectin exerts a potent protective activity on microglia cells and strongly support further investigations aimed at exploring its therapeutic role in *in vivo* models of neurodegenerative disorders.

## 1. Introduction

Neurological disorders (NDs), including neurodegenerative diseases, are one of the most impactful conditions afflicting contemporary society wellness, being the second cause of death worldwide (Feigin et al., 2020). The growth of the global population, aging and unhealthy and industrialized lifestyle are leading to an exponential increase in the burden of NDs. Moreover, NDs emerge as multifactorial diseases, sharing pathogenic pathway that includes mitochondrial dysfunction, oxidative stress, misfolded protein aggregation, and neuroinflammation. Consequently, despite decades of clinical and basic research, most therapeutic strategies designed to manage degenerative NDs are only palliative and often associated with a broad spectrum of side effects (Passeri et al., 2022).

Microglia cells, descending from the myeloid lineage, are considered the resident immune cells in the brain. In a physiological context, resting microglia shows a ramified body, which allows to oversee neuronal status and the biochemical balance of the surrounding area, by producing neurotrophic factors, executing the synaptic pruning, and exerting a phagocytic activity of cellular debris. Under pathological chronic triggers, microglia changes morphology and activates a cascade of events which leads to the failure of the brain homeostasis surveillance, the activation of inflammatory response, and enhances the neurodegeneration process (Heneka et al., 2015). Neuroinflammation and neurodegeneration are sustained by the release of pro-inflammatory cytokines, including interleukin (IL)-1β and IL-6, chemokines, small-molecule messengers such as prostaglandins and nitric oxide (NO), and the activation of molecular pathways involving nuclear factor kappa-light-chain-enhancer of activated B cells (NF-κB) and reactive oxygen species (ROS) (Leng and Edison, 2021). In particular, the abnormal amount of ROS and the establishment of oxidative stress create a synergy with neuroinflammation, and the two processes are intrinsically related phenomena leading to neurodegeneration (Teleanu et al., 2022) (Reynolds et al., 2007). For all these reasons, microglia cells represent an important target for the development of therapeutic strategies, and the full understanding of their response under basal and pathological conditions is essential in the context of preclinical and clinical neuroscience research (Dello Russo et al., 2018).

So far synthetic drugs failed in preventing or treating neurodegenerative diseases. Plentiful research efforts, thus, have been focused on possible use of natural and bioactive compounds derived from plants, fruits, and seeds exerting antioxidant and anti-inflammatory actions (Chen et al., 2022) (Stacchiotti and Corsetti, 2020) (Di Liberto and Mudo, 2022). A common neuroprotective mechanism for widely different natural compounds derives from their ability to scavenge the excess of free radicals generated in oxidative and neurotoxin-induced processes in the brain nerve cells (Wasik and Antkiewicz-Michaluk, 2017). In the context of these researches, we have lately shown the powerful neuroprotective, antioxidant and mitoprotective activity of lemon (Nuzzo et al., 2021a) and grapefruit (Nuzzo et al., 2021b) IntegroPectin in neuronal-like cells. “IntegroPectin” is the name of a new family of *Citrus* pectins obtained via hydrodynamic cavitation of *Citrus* fruit processing industrial biowaste rich in adsorbed flavonoids (Scurria et al., 2021) and terpenes (Scurria et al., 2020) characterized by a broad biological activity. In the first *in vivo* experiments, grapefruit IntegroPectin recently showed anti-ischemic cardioprotective activity, significantly higher than pure naringenin, namely the bioactive aglycone of naringin (Flori et al., 2022).

In this study, we have investigated the anti-apoptotic and anti-inflammatory properties of grapefruit IntegroPectin on human microglial clone 3 (HMC3) cell line (resulting from the transfection of the SV40 T antigen in primary human microglial cultures, derived from 8- to 10-week old embryos), focusing on the modulation of specific intracellular signalling cascades. These cells retain most of the morphological and phenotypical characteristics of the primary source: they look like globular or elongated cells with thick processes and dark cytoplasmic granulation; they are positive for microglial and monocyte markers such as Cluster of Differentation 68 (CD 68) and Ionized calcium-binding Adapter molecule 1 (IBA1); and, they are able to release pro-inflammatory factors, such as IL-6, even under basal condition. HMC3 cells can be activated by different stimuli, including pro-oxidant and neuroinflammatory activators (Dello Russo et al., 2018). In the present study, HMC3 cells were challenged with *tert*-butyl hydroperoxide (TBH), a relatively stable hydroperoxide inducing oxidative stress (Pias and Aw, 2002), in the presence of grapefruit IntegroPectin, and the response was measured.

## 2. Materials and methods

### 2.1 Solubilization of IntegroPectin

Grapefruit IntegroPectin, obtained as previously described from grapefruit processing industrial waste (Scurria et al., 2021) kindly donated by Campisi Citrus (Siracusa, Italy), was solubilized at the concentration of 10 mg/mL in cell culture medium. The solution was filtered using a 0.45 μm filter (Sartorius) and stored at 4 °C.

### 2.2 Cell cultures and treatments

HMC3 cells, generously donated by Prof. M. Sortino (University of Catania), were cultured in T25 tissue culture flasks in a humidified atmosphere of 95% air and 5% CO_2_ at 37 °C. Cells were used at the passage number range 8-14. The growth medium was composed by MEM (Minimum Essential Medium) – w/Earle’s Salts, supplemented with 10% fetal bovine serum (FBS), 100 U/mL penicillin, 100 U/mL streptomycin, 2 mM L-glutamine and non-essential amino acids. The cell culture medium was changed once a week and the cells were sub-cultured once they reached 90% confluence. All treatments were performed at least 48 h after plating. Based on the experimental groups, the cells received the following treatments: IntegroPectin 1 mg/mL, 0.1 mg/mL and 0.01 mg/mL for 24 h in dose-effect experiments; IntegroPectin 1 mg/mL for 24 h, 48 h and 72 h in time-course experiments; TBH 200 μM, 100 μM and 50 μM for 24 h in dose-response experiments; a combination of IntegroPectin (1 mg/mL) and TBH (200 μM), with IntegroPectin administered immediately before (preventive co-treatment) TBH treatment (24 h); in experiments requiring treatment with PD98059 (Tocris 1213), an inhibitor of mitogen-activated protein kinase (MAPK), and LY294002 (440202 Sigma-Aldrich), a phosphoinositide 3-kinase (PI3K)/Akt inhibitor (Jiang et al., 2010), the compounds (30 μM and 10 μM, respectively) were administered 1 h before IntegroPectin and TBH exposure. The dose of the two inhibitors were selected based on manufacturer instructions and the broad existing literature (Lu et al., 2022) (Battaglia et al., 2009). For experiments aimed at assessing the modulation of mRNA expression and acute intracellular pathway modulation, cells were treated for 4 h with IntegroPectin 1 mg/mL. In all experiments, the control group was treated with an equal volume of cell medium.

### 2.3 Cell viability by MTT assay

Cells were grown at a density of 2.5 × 10^4^ cells/well in 96-well plates in a final volume of 100 μL/well. Cellular metabolic activity, an indicator of cell viability, was assessed by measuring the intracellular reduction of tetrazolium salt (MTT, 0.5 mg/mL) to purple formazan granules by mitochondrial succinate dehydrogenase expressed in metabolically active cells, after 3 h incubation at 37 °C. Absorbance was measured at 570 nm with background subtraction after solubilizing MTT–formazan product with dimethyl sulfoxide (DMSO), 100 μL/well. Cell viability was expressed as percentage with the control group set to 100.

### 2.4 Cell viability and nuclear morphology by 4′,6-diamidino-2-phenylindole (DAPI) staining

For analysis of cell nuclei morphology, cells were grown at a density of 2 × 10^5^ cells/well on coverslips placed in a 24-well plate in a final volume of 400 μL/well. Cells were fixed with 4% formaldehyde solution for 15 min at room temperature and, after two washings with phosphate buffered saline (PBS), nuclei were counterstained with the fluorescent stain (DAPI). After one washing with PBS, the coverslips were mounted on slides, and the cellular images were obtained using a fluorescence microscope (DMRBE, Leica Microsystems GmbH, Germany), equipped with digital video camera (Spot-RT Slider, Diagnostic Instruments, Mi, USA).

### 2.5 Quantification of Reactive Oxygen Species (ROS) by dichlorofluorescein diacetate (DCFH-DA) assay

HMC3 cells were placed at a density of 2.5 x 10^4^ cells/well in 96-well plates in a final volume of 100 μL/well. After exposure to the indicated treatments, cells were incubated in the dark for 10 min at room temperature with DCFH-DA (1 mM), washed with PBS and fluorescence intensity was measured by using the microplate reader GloMax fluorimeter (Promega Corporation, Madison, WI, USA) at the excitation wavelength of 475 nm and emission wavelength 555 nm. Results were expressed as percentage with the control group set at 100.

### 2.6 Western Blotting analysis

To assess Caspase-3, phosphorylated (p)-Extracellular signal-regulated kinase (ERK)1/2, p-Akt and p-NF-kB proteins expression levels, HMC3 cells were placed at a density of 2.5 x 10^4^ cells/well in 96-wells plates in a final volume of 100 μL/well. At the end of treatment, for each experimental condition, cells were mechanically detached and the cell content of 4 wells was pooled into one sample. The cell pellet was homogenized in cold radioimmunoprecipitation assay (RIPA) buffer (50 mM Tris, pH 7.4, 150 mM NaCl, 1% Triton, SDS 0.1%), supplemented with protease inhibitor cocktail (Sigma-Aldrich P8340) and phosphatase inhibitor cocktail (Sigma-Aldrich P5726) and sonicated (30 pulsations/min). Proteins were quantified by the Lowry method (Lowry et al., 1951). Western blotting procedure was performed as previously described (Scordino et al., 2023). Protein samples (30 μg per lane) and molecular weight marker (PageRuler Plus Prestained Protein Ladder, 26619 ThermoFisher Scientific) were run on 8% polyacrylamide gel and electrophoretically transferred onto nitrocellulose membrane (RPN303E, Hybond-C-extra, GE Healthcare Europe). The membranes were incubated at 4 °C under gentle shaking for 1 h in blocking buffer (1× TBS, 0.1% Tween−20, 5% w/v nonfat dry milk), followed by overnight incubation with specific antibodies diluted in blocking buffer. For the detection of Caspase-3, p-ERK1/2, p-Akt, and p-NF-kB levels the following antibodies were used: mouse anti-Caspase-3 (1:400, sc-56053 Santa Cruz Biotechnology), rabbit anti-p-ERK1/2 (1:1000, 9101 Cell Signaling Technology), rabbit anti-p-Akt (1:1000, 4060 Cell Signaling Technology), mouse p-NF-kB (1:500, sc-136548 Santa Cruz Biotechnology). The day after, membranes were washed three times for 10 min with TBS/T and incubated for 1 h with goat anti-mouse IgG-HRP (sc-2005 Santa Cruz Biotechnology) or mouse anti-rabbit IgG-HRP (sc-2357 Santa Cruz Biotechnology) horseradish peroxidase conjugated diluted 1:10000. After three washings with TBS-T, immunocomplexes were visualized with chemiluminescence reagent (SuperSignal West Pico Plus, ThermoFisher Scientific). The membranes were exposed to autoradiography film (Amersham Hyperfylm ECL, 28-9068-36). Chemiluminescent signal was visualized and fixed in Kodak D19 developer and fixer (1900984 and 1902485, Kodak GBX). A sample of chemiluminescent membranes was developed by using iBright FL1000 Imaging System (ThermoFisher Scientific). For western blotting normalization, membranes were washed and exposed to horseradish peroxidase-conjugated β-Actin primary antibody (sc-47778 Santa Cruz Biotechnology), diluted 1:10000, for 1 h. The densitometric evaluation of bands was performed by measuring the optical density (O.D.) using NIH ImageJ software with results being expressed as arbitrary units.

### 2.7 Cytofluorimetry analysis for Annexin V/Propidium Iodide (PI) apoptosis assay

HMC3 cells were plated at a density of 2.5 × 10^4^ cells/well in 96-well plates in a final volume of 100 μL/well. At the end of treatment cells were mechanically detached and centrifuged at 1200 rpm for 5 min at room temperature. After discarding the supernatant, samples were resuspended in cold PBS 1X -/-(no calcium, no magnesium) (Gibco, Thermo Fisher Scientific) and centrifuged. Cells were resuspended at 1×10^5^ cells/ in 100 μl of Annexin-binding buffer 1X (Invitrogen, Thermo Fisher Scientific) and 5 μL of Alexa Fluor 647 Annexin V (Invitrogen, Thermo Fisher Scientific) diluted 1:10 with Annexin V binding buffer 1X were added to each 100 μl of cell suspension. An unstained control was prepared. All the samples were incubated at room temperature for 15 minutes in the dark. After incubation, cold PBS was added and cells were centrifuged, the supernatants discarded to remove any excess of unbound Annexin V, resuspended in 200 μl of Annexin-binding buffer 1X and kept on ice. PI (1 μl/100 μl) was added 5 minutes before starting the reading on the flow cytometer CytoFlex S (Beckman Coulter, Brea, CA, USA). Results were shown and analyzed through CytExpert flow cytometry analysis software.

### 2.8 RNA isolation and reverse transcription PCR

HMC3 cells were cultured at a density of 2.5 x 10^4^ cells/well in 96-wells plates in a final volume of 100 μL/well. At the end of treatment, for each experimental condition, cells were mechanically detached and the cell content of 7 wells was combined into one sample. The RNA was isolated from the cell pellet using a Qiagen RNeasy mini kit (74104, Qiagen, Hilden, Germany) following the manufacturer’s instructions. The total RNA concentration was detected using a Thermo Scientific MultiskanGO Microplate Spectrophotometer and 1 μg was immediately retro-transcribed using a mixture containing: 5× first strand buffer (18080-044, Invitrogen,), 2.5 μM of random hexamers (N112470, Roche), 2.5 mM of dithiothreitol (DTT) (18080-044, Invitrogen), 0.5 mM of dNTPs mix (18427-013, Invitrogen), 40U of RNAse inhibitor (N8080119, Invitrogen), 200 U of Superscript III Reverse Transcriptase (18080-044, Invitrogen). Reaction mixtures (20 μL) were incubated for 2 h at 50 °C and then for 15 min at 70 °C.

### 2.9 Real-time PCR

Quantitative gene expression analysis of IL-6, IL-1β, Inducible nitric oxide synthase (iNOS) was performed using SYBR Green Real-time PCR. The reaction was carried out in a total volume of 20μl containing: Power Track SYBR Green Master mix (A46109, AppliedBiosystem), 2 μL of cDNA, and 0.6 μM of primers mix (Forward (5’ to 3’): IL6 CACTGGTCTTTTGGAGTTTGAG; IL1β GCCAGTGAAATGATGGCTTATT; iNOS GACTTTCCAAGACACACTTCAC; Reverse (5’ to 3’) IL6 GGACTTTTGTACTCATCTGCAC; IL1β AGGAGCACTTCATCTGTTTAGG; iNOS TTCGATAGCTTGAGGTAGAAGC). RT-PCR was performed in 48-well plates using the Step-One Real-Time PCR System (Applied Biosystems). Relative changes in gene expression between control and treated samples were determined using the 2 (-Delta Delta CT) method. Levels of the target transcript were normalized to β-actin levels (Forward primer: TCCCTTGCCATCCTAAAAGCCACCC; Reverse primer: CTGGGCCATTCTCCTTAGAGAGAAG). Final values were expressed as fold change.

### 2.10 Statistical analysis

All the quantitave analyses were carried out in a blind manner by two independent experimenters unware of sample identity. Statistical data analysis was performed using GraphPad Prism 9.0.2 software (GraphPad Software, La Jolla, CA, USA). Normal distribution of data was assessed by Shapiro-Wilk test. Statistical evaluations were performed by one-way ANOVA, followed by Tukey Post-Hoc test. t-test was used for the statistical comparison between the means of two groups. The relative results were presented as mean ± SE of at least three independent experiments. Differences in *p*-value less than 0.05 were considered statistically significant.

## 3. Results

### 3.1 Effects of grapefruit IntegroPectin on HMC3 cell viability

We first evaluated the viability of HMC3 cells after treatment with grapefruit IntegroPectin. Results from dose-response experiments indicate no changes in cell viability following cell exposure for 24 h to three different doses of IntegroPectin: 0.01 mg/mL, 0.1 mg/mL and 1 mg/mL (Figure 1A). Next, we carried out a time-course experiment by treating cells with the highest dose of IntegroPectin (1 mg/mL). Results in Figure 1B show that IntegroPectin does not induce any significant changes in cell viability, even when the treatment is prolonged up to 72 h. Based on these data, IntegroPectin 1 mg/mL was used in all the subsequent experiments.

**Figure 1.**
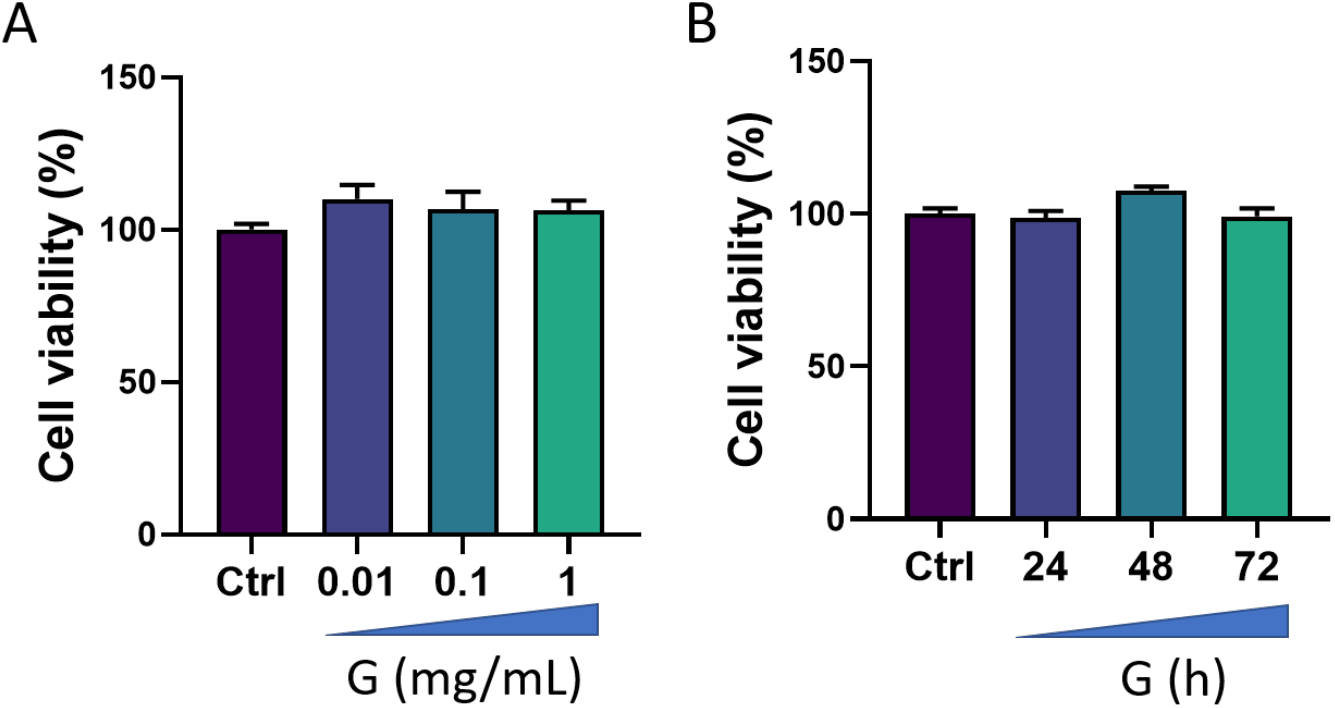
Effects of grapefruit IntegroPectin (G) on HMC3 cell viability. A) Dose–effect of G treatment (24 h) on cell viability, evaluated by MTT assay (n=3 experiments, total n=39):. B) Time-course of G treatment (1 mg/mL) effects on cell viability, assessed by MTT assay (n=3 experiments, total n=31).

### 3.2 Protective and antioxidant effects of grapefruit IntegroPectin in HMC3 cells treated with TBH

We then tested the ability of IntegroPectin to recover cell viability impaired by oxidizing agent (TBH) treatment. HMC3 exposure for 24 h to TBH leads to a dose-dependent decrease in cell viability (Figure 2A). We selected TBH highest dose (200 μM), which is able to induce a significant decrease in cell viability, for all the subsequent experiments. To evaluate the protective effect of IntegroPectin, HMC3 cells were treated with IntegroPectin (1 mg/mL) immediately before TBH exposure. As shown in Figure 2B, IntegroPectin treatment fully prevents the cell death induced by TBH treatment.

**Figure 2.**
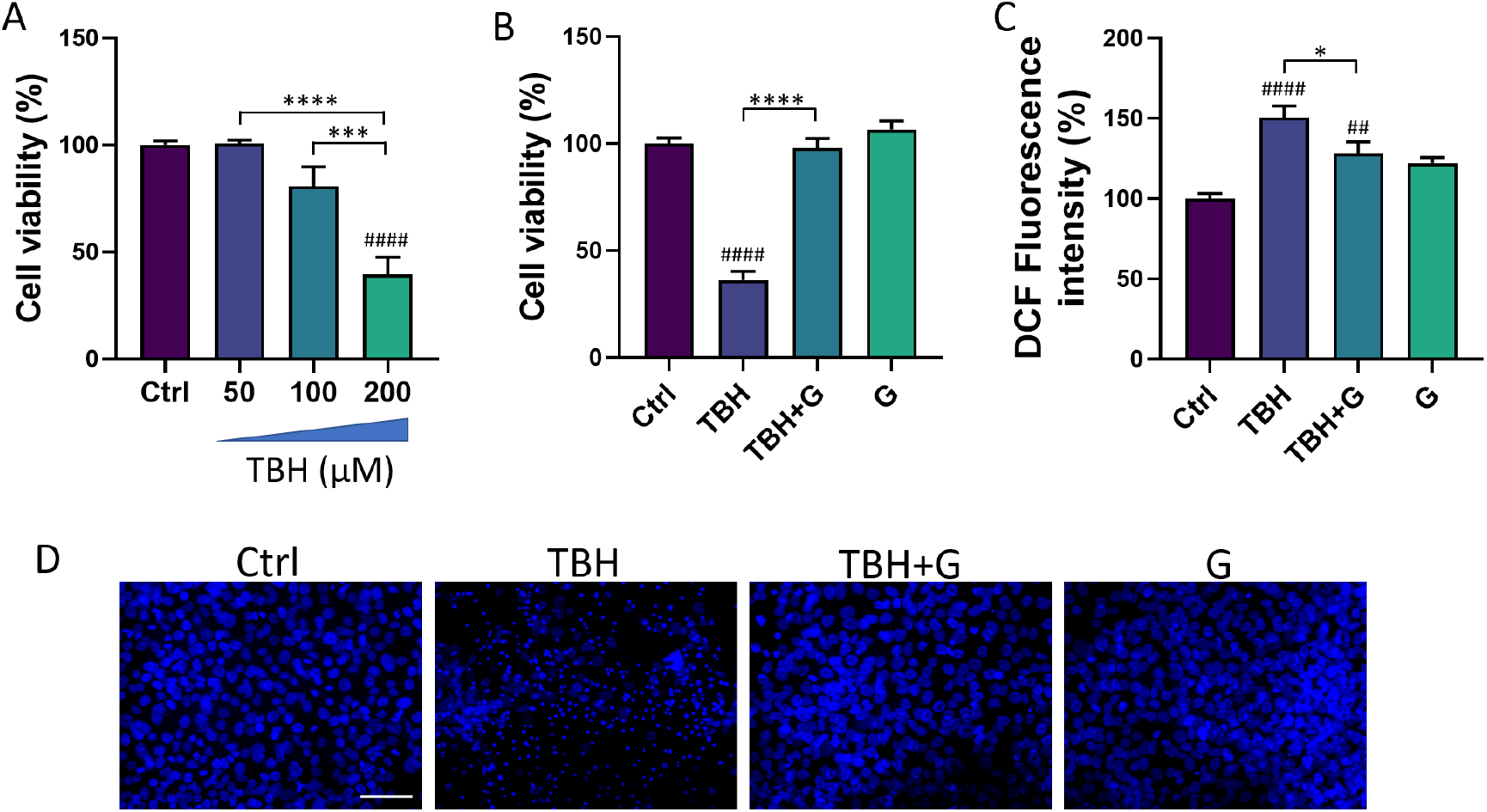
Protective and antioxidant effects of grapefruit IntegroPectin (G). A) Dose–response effects of TBH treatment (24 h) on HMC3 cell viability, evaluated by MTT assay (n=3 experiments, total n=27). B) Quantification of cell viability by MTT test in untreated (Ctrl) cells, cells treated with TBH (200 μM, 24 h), TBH (200 μM, 24 h) + G (1 mg/mL, 24 h), and G alone (1 mg/mL, 24 h) (n=3 experiments, total n=44). C) DCF fluorescence intensity quantification, an index of intracellular ROS generation, in Ctrl cells, cells treated with TBH (200 μM, 24 h), TBH (200 μM, 24 h) + G (1 mg/mL, 24 h), and G alone (1 mg/mL, 24 h). (n=3 experiments, total n=47) D) Representative pictures of DAPI nuclear staining in Ctrl cells, cells treated with TBH (200 μM, 24 h), TBH (200 μM, 24 h) + G (1 mg/mL, 24 h), and G alone (1 mg/mL, 24 h). Tukey test: ## p < 0.01, #### p < 0.0001 as compared with Ctrl group. *** p<0.05, *** p < 0.001, **** p < 0.0001. Scale bar: 100μm.

The impact of IntegroPectin on TBH-induced oxidative stress and ROS generation was assessed by DCFH-DA fluorescence intensity assay. Fluorescence intensity measurement, an index of intracellular ROS levels, shows that treatment of HMC3 cells with grapefruit IntegroPectin significantly reduces intracellular ROS generation, boosted by TBH exposure (Figure 2C).

Moreover, the protective effects exerted by IntegroPectin treatment on cell viability following TBH exposure were further confirmed by qualitatively comparing the morphology of DAPI stained-cell nuclei. Bright fluorescent nuclei of TBH-treated cells appear smaller, fragmented and condensed, as compared to control nuclei, while treatment with grapefruit Integropectin prevents this morphological change (Figure 2D).

### 3.3 Anti-apoptotic effects of grapefruit IntegroPectin in HMC3 cells treated with TBH

Cell death induced by TBH exposure is mainly mediated by the activation of the apoptotic process (Pias and Aw, 2002). In order to study the ability of grapefruit IntegroPectin to inhibit apoptosis induced by TBH treatment, cellular apoptosis/necrosis was evaluated by flow cytometry quantification of Annexin V binding and PI uptake. In Figure 3A, values within the representative density plots indicate the percentage of viable Annexin V^-^ PI^-^ cells, intermediate apoptotic Annexin V^+^ PI^-^ cells, necrotic Annexin V^-^ PI^+^ cells, and late apoptotic Annexin V^+^ PI^+^ cells. Plots show that cells in the untreated (Ctrl) sample retain high viability (Annexin V^-^ PI^-^, 95.5%) with only a small percentage of Annexin V^+^ PI^-^ cells (4.3%) undergoing intermediate apoptosis. The oxidizing action of TBH affects the viability of the sample (Annexin V^-^ PI^-^ 36%) leading to a higher percentage of Annexin V^+^ PI^-^ cells (63%) indicative of intermediate apoptosis. In a co-treatment with TBH, IntegroPectin exerts a protective action as the sample yield a high percentage of Annexin V^-^ PI^-^ cells (95%) and a small percentage (4.5%) of Annexin V^+^ PI^-^ cells, thus preventing oxidative stress-induced apoptosis. Treatment with IntegroPectin alone does not induce any change in cell viability and in the percentage of Annexin V^-^ PI^-^ cells (97.2%) and Annexin V^+^PI^-^ cells (2.5%), as compared to the ctrl group. No significant percentages of necrotic cells (Annexin V^-^ PI^+^) or secondary necrotic cells (Annexin V^+^ PI^+^) show up in both density plots. Histograms in Figure 3B and Figure 3C gather the cumulative percentages of viable Annexin V^-^ PI^-^ cells and intermediate apoptotic Annexin V^+^ PI^-^ cells in different experiments. In order to further characterize the anti-apoptotic role of grapefruit IntegroPectin, the protein levels of Caspase-3, a thiol protease that acts as a major effector caspase involved in the execution phase of apoptosis (Brentnall et al., 2013), were explored. Treatment with the oxidizing agent alone (TBH 200 μM) induces a significant increase in the cleaved forms of Caspase-3 (25 kDa and 17 kDa), while grapefruit IntegroPectin is able to counteract this effect (Figure 3D).

**Figure 3.**
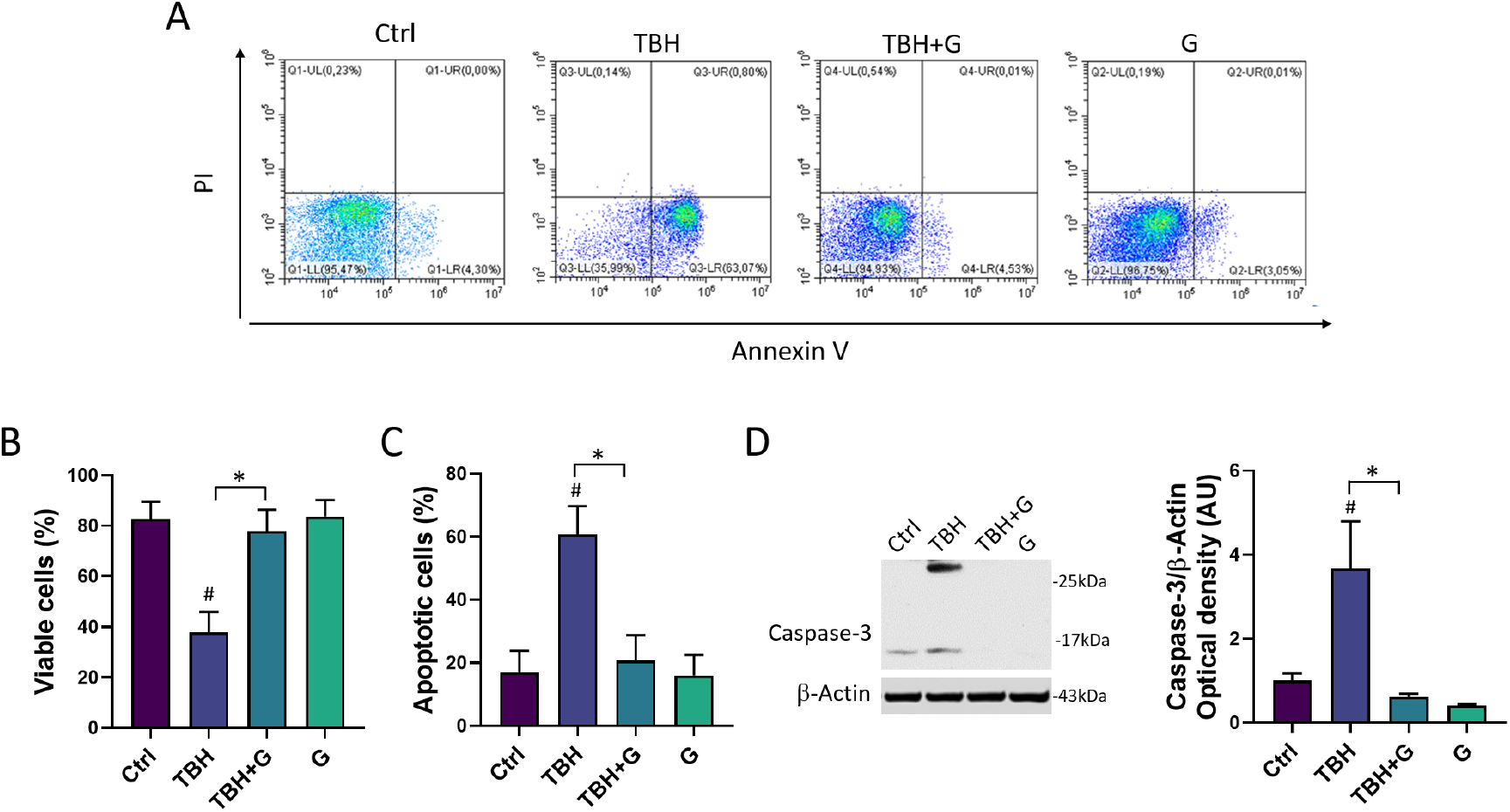
Anti-apoptotic effects of grapefruit IntegroPectin. A) Representative plot indicating the percentage of Annexin V-PI-cells (on the lower left quadrant), Annexin V+ PI-cells (on the lower right quadrant), PI+ Annexin V-cells (on the upper left quadrant) and PI+ Annexin V+ cells (on the upper right quadrant) of different conditions: untreated sample (Ctrl), sample treated with TBH (200 μM, 24 h), sample co-treated with TBH (200 μM, 24 h) and IntegroPectin (G) (1 mg/mL, 24 h), sample treated with IntegroPectin (G) alone (1 mg/mL, 24 h). b) Histogram showing the cumulative Annexin V-PI-cell (viable cells) percentages from all the experiments (n=3 experiments, total n=12). c) Histogram showing the cumulative AnnexinV+ PI-cells percentages (apoptotic cells) from all the experiments (n=3 experiments, total n=12). d) Representative images of Caspase-3 and β-actin Western blotting bands and histogram of Caspase-3 normalized to β-actin Optical density in Ctrl cells, cells treated with TBH (200 μM, 24 h), TBH (200 μM, 24 h) + G (1 mg/mL, 24 h), and G alone (1 mg/mL, 24 h) (n=4 experiments, total n=15). Tukey test: # p < 0,05 as compared to Ctrl group; * p < 0,05. AU (Arbitrary Units).

### 3.4 Modulation of MAPK/ERK and PI3K/Akt pathways by grapefruit IntegroPectin

In order to characterize the molecular mechanisms underlying grapefruit IntegroPectin protective response, we explored the modulation of two major cell survival pathways: MAPK/ERK and PI3K/Akt. As shown in Figure 4A, HMC3 cells treated with TBH (200 μM) show a significant down-regulation of phosphorylated (p)-ERK1/2 levels, which is fully counteracted by grapefruit IntegroPectin (G, 1 mg/mL) co-treatment. Surprisingly, grapefruit IntegroPectin alone is able to significantly enhance p-Erk protein levels, as compared to untreated (Ctrl) cells. In order to explore the specific involvement of MAPK/ERK pathway in IntegroPectin protective function, HMC3 cells received a treatment with PD98059 (30 μM), a selective ERK1/2 signaling inhibitor, 1 h before grapefruit IntegroPectin (1 mg/mL) and TBH (200 μM) exposure (24 h). Treatment with PD98059 slightly reduces IntegroPectin protective effects against oxidative stress-induced cell death (Figure 4B), suggesting a minor involvement of MAPK/ERK signaling. In addition to p-ERK1/2 levels, IntegroPectin (1 mg/mL) co-treatment is able to fully recover phosphorylated (p)-Akt protein levels, down-regulated by TBH exposure (200 μM, 24 h) (Figure 4C). Interestingly, when HMC3 cells were treated with LY294002 (10 μM), a selective PI3K/Akt inhibitor, 1 h before exposure to grapefruit IntegroPectin (G, 1 mg/mL) and TBH (200 μM, 24 h), the IntegroPectin protective activity completely vanished (Figure 4D), suggesting the major involvement of PI3K/Akt signaling in this effect.

**Figure 4.**
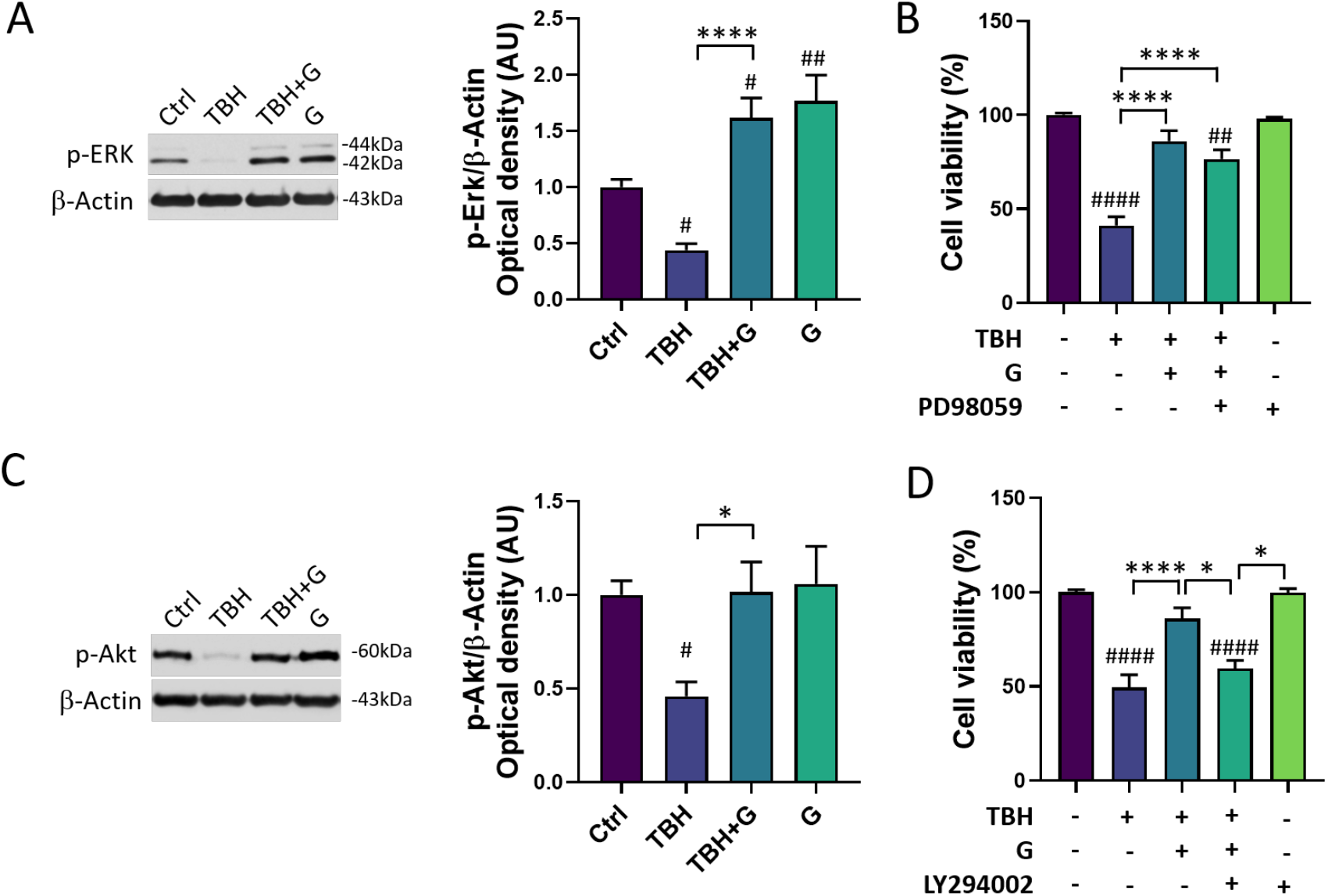
Modulation of MAPK/ERK and PI3K/Akt pathways by grapefruit IntegroPectin. A) Representative images of phosphorylated (p)-ERK1/2 and β-Actin Western blotting bands and histogram of p-ERK1/2 normalized to β-Actin Optical density in untreated (Ctrl) cells, cells treated with TBH (200 μM, 24 h), TBH (200 μM, 24 h) + IntegroPectin (G) (1 mg/mL, 24 h), and G alone (1 mg/mL, 24 h) (n=4 experiments, total n=35). B) Quantification of cell viability by MTT test in Ctrl cells, cells treated with TBH (200 μM, 24 h), TBH (200 μM, 24 h) + G (1 mg/mL, 24 h), TBH (200 μM, 24 h) + G (1 mg/mL, 24 h) + PD98059 (30 μM), and PD98059 (30 μM) only. PD98059 was administered 1 h before grapefruit IntegroPectin and TBH exposure. (n=5 experiments, total n=71). C) Representative images of phosphorylated (p)-Akt and β-Actin Western blotting bands and histogram of p-Akt normalized to β-Actin Optical density in Ctrl cells, cells treated with TBH (200 μM, 24 h), TBH (200 μM, 24 h) + G (1 mg/mL, 24 h), and G (1 mg/mL, 24 h) alone (n=5 experiments, total n=36). D) Quantification of cell viability by MTT test in Ctrl cells, cells treated with TBH (200 μM, 24 h), TBH (200 μM, 24 h) + G (1 mg/mL, 24 h 10), TBH (200 μM, 24 h) + G (1 mg/mL, 24 h) + LY294002 (10 μM), and LY294002 (10 μM) only. LY294002 was administered 1 h before grapefruit IntegroPectin and TBH exposure (n=3 experiments, total n=59) Tukey test: # p < 0,05, ## p < 0,01, #### p < 0,0001 as compared to Ctrl group; * p < 0,05, **** p < 0,0001. AU (Arbitrary Units).

### 3.5 Modulation of neuroinflammatory pathways by grapefruit IntegroPectin

In order to assess the potential anti-inflammatory power of grapefruit IntegroPectin, we investigated the mRNA expression of three major genes involved in the modulation of inflammatory response in microglia cells: the main pro-inflammatory cytokines IL-6 and IL-1β, and the major downstream mediator of inflammation iNOS (Zamora et al., 2000). While IntegroPectin (G) treatment does not modulate the expression of IL-6 (Figure 5A) and IL-1β (Figure 5B), the bioactive molecule is able to significantly reduce the expression of iNOS, as compared to untreated (ctrl) cells (Figure 5C).

**Figure 5.**
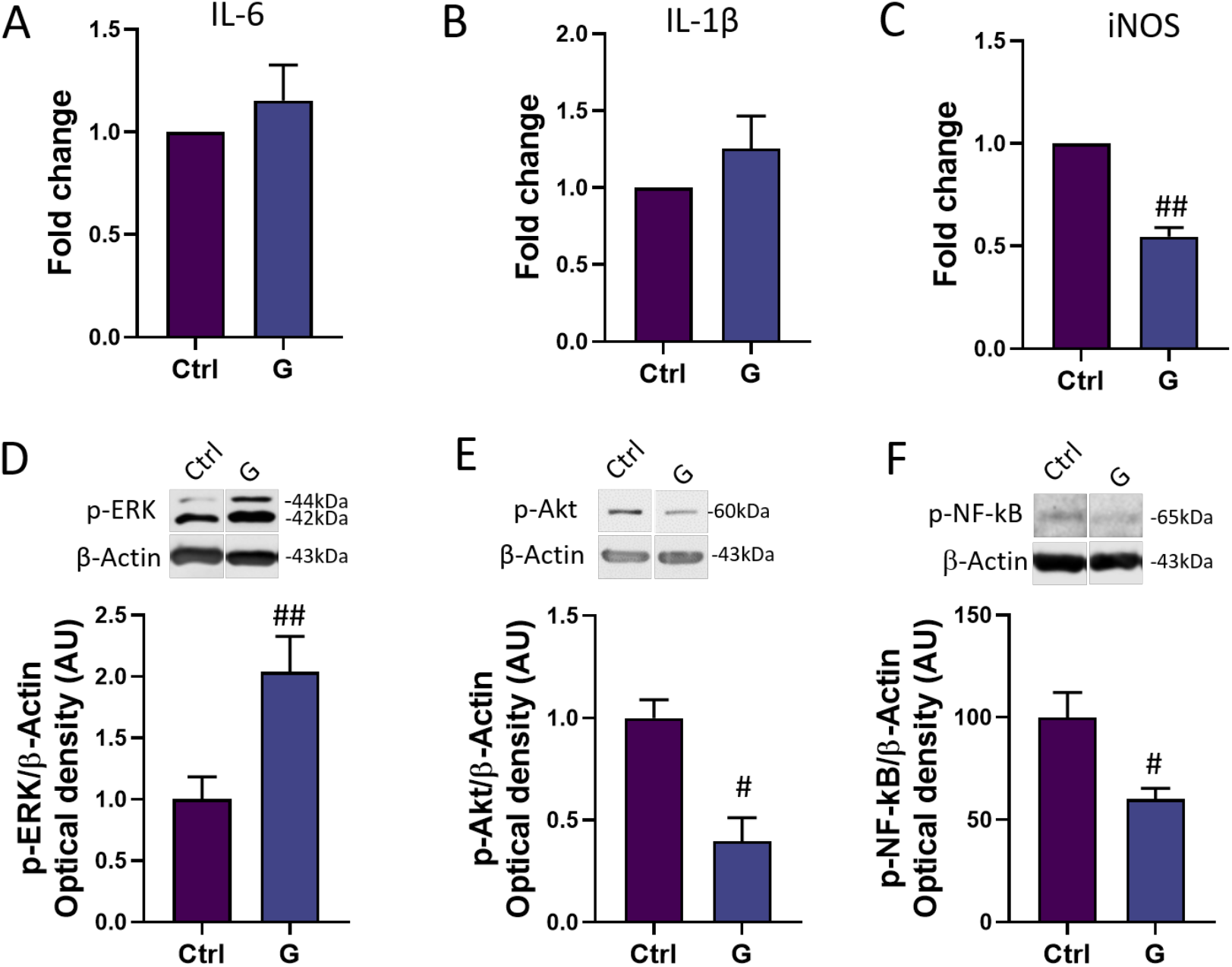
Modulation of inflammatory mediators and pathways by grapefruit IntegroPectin. RT-PCR of IL-6 (n=3 experiments, total n=6) (A), IL-1β (n=3 experiments, total n=6) (B), and iNOS (n=3 experiments, total n=6) (C) mRNA levels in untreated (Ctrl) cells and cells treated with IntegroPectin (G) (1 mg/mL, 4 h). Representative images of phosphorylated (p)-ERK1/2 (D), p-Akt (E), p-NF-kB (F) and β-Actin Western blotting bands and histogram of p-ERK1/2 (n=3 experiments, total n=18) (D), p-Akt (n=3 experiments, total n=6) (E) and p-NF-kB (n=5 experiments, total n=10) (F) normalized to β-Actin Optical density in Ctrl cells and cells treated with IntegroPectin (G) (1 mg/mL, 4 h). t-test: # p < 0,05, ## p < 0,01, as compared to Ctrl group. AU (Arbitrary Units).

We also explored the time-related activation of three important intracellular modulators of the neuroinflammatory response: ERK1/2, Akt and NF-kB. Interestingly, a short-time (4 h) IntegroPectin (G) treatment leads to a significant increase in p-ERK1/2 (Figure 5D) levels and a parallel decrease in p-Akt (Figure 5E) and p-NF-kB (Figure 5F) levels, as compared to Ctrl cells.

## 4. Discussion

Grapefruit IntegroPectin is a highly water-soluble *Citrus* pectin characterized by the preservation of the Rhamnogalacturonan-I (RG-I) backbone and endowed with large amounts of bioactive molecules, including flavonoids (Scurria et al., 2021) and terpenes (Scurria et al., 2020) well known for their robust anti-oxidant and anti-inflammatory properties. In line with the results of the present study, previous data have described the neuroprotective (Nuzzo et al., 2021b) effects of grapefruit IntegroPectin in neuronal-like cells and its anti-ischemic cardioprotective (Flori et al., 2022) activity, suggesting the activation of multi-target beneficial effects by this biopolymer.

So far, the effects of IntegroPectin on microglia cells and the modulation of neuroinflammatory response was unexplored. By a combination of several experimental approaches, including nuclei morphology, Annexin V binding-PI uptake, and quantification of Caspase-3 levels, a protein crucial for apoptotic chromatin condensation and DNA fragmentation (Porter and Janicke, 1999), we observed that grapefruit IntegroPectin is able to counteract TBH-induced apoptosis in human microglial HMC3 cell line. We can speculate that the scavenger activity of grapefruit IntegroPectin against ROS blocks the cross-linked extrinsic and intrinsic apoptotic pathways, in the end re-establishing the mitochondrial homeostasis, as recently discovered in SH-SY5Y cells using both grapefruit (Nuzzo et al., 2021b) and lemon (Nuzzo et al., 2021a) IntegroPectin. Interestingly, while grapefruit IntegroPectin treatment in SH-SY5Y cells produced a decrease in the proliferation rate and a cell cycle arrest at the G2/M phase, the same treatment does not affect cell viability and cell proliferation in HMC3 cells, suggesting the activation by IntegroPectin of cell-specific effects.

Microglia exposure to TBH leads to a decrease in PI3K/Akt and MAPK/ERK signaling activation. Both effects are counteracted by IntegroPectin treatment. PI3K/Akt and MAPK/ERK pathways have been widely described as pro-survival pathways (Ong et al., 2016), essential for cell protection against a plethora of noxious stimuli, including oxidative stress. Flavanones, for example, exert their neuroprotective activity by modulating the apoptosis process through Akt and ERK1/2 signaling pathways (Vauzour et al., 2007). Moreover, these two intracellular pathways are also involved in the neuroprotective effects activated by ginseng pectin treatment in cortical neuron cells (Fan et al., 2012). Interestingly, using specific inhibitors, here we disclose a prominent role of PI3K/Akt cascade in guaranteeing cell survival following cell exposure to TBH.

More in general, both MAPK/ERK and PI3K/Akt pathways represent a point of convergence of several cellular processes, including inflammation. Experimental evidence indicates that the PI3K/Akt pathway is required for lipopolysaccharide (LPS) activation of microglial cells (Saponaro et al., 2012), while other studies suggest that PI3K/Akt activation contributes to the anti-inflammatory effects exerted by several biomolecules (Sun et al., 2016), thus outlining a complex scenario of both protective and harmful responses orchestrated by this intracellular cascade. Similar results were obtained exploring the role of MAPK/ERK pathway in neuroinflammation modulation: an increase in ERK activation has been associated with the suppression of inflammatory genes in endothelial cells (Maeng et al., 2006), and with the neuroinflammatory effects of Astragaloside IV in LPS induced microglial cells (Harrison et al., 2018). Conversely, ERK is a critical regulator of Interferon (IFN)γ-mediated pro-inflammatory activation of microglia (Chen et al., 2021), and several anti-inflammatory drugs act through MAPK/ERK down-regulation (Lim et al., 2018). These pieces of evidence suggest that several experimental variables, including cell model, duration and typology of stimuli, may contribute to the state of activation of these pathways in the context of microglia physiology.

In our cell model, a short (4 h) treatment with IntegroPectin is able to induce the up-regulation of MAPK/ERK activation, together with a down-regulation of the PI3K/Akt cascade. Interestingly, we observed that IntegroPectin treatment of microglia cells leads also to the decrease in NF-kB activation, a transcriptional master regulator of the inflammatory and the apoptotic response (Tak and Firestein, 2001). The canonical NF-κB is activated in response to several external stimuli involved in inflammation, immune response, cell proliferation, differentiation, and survival, through phosphorylation and subsequent degradation of the inhibitory IκB protein. On the other hand, the non-canonical NF-κB is selectively activated by TNF superfamily receptors which lead to the activation of NF-κB-inducing kinase (NIK), NIK-mediated p100 phosphorylation and nuclear translocation of NF-κB (Yu et al., 2020). We can hypothesize that the basal activation of NF-κB pathway in HMC3 cells is likely dependent on the PI3k-Akt cascade, which regulates the transcriptional activity of NF-κB by inducing the phosphorylation and the subsequent degradation of inhibitor of κB (IκB) (Kane et al., 1999). By modulating Akt activation, grapefruit IntegroPectin leads to the subsequent downregulation of NF-kB activity and inflammatory response. In line with our results, recent studies described the beneficial activity of a modified citrus pectin in the inhibition of the NLRP3 inflammasome and NF-kB pathway activation in microglia during cerebral ischemia (Cui et al., 2022).

We also investigated the expression of a panel genes typically associated with the pro-inflammatory response and the activation of the PI3K-NF-kB cascade (Natarajan et al., 2018). Surprisingly, while IntegroPectin treatment does not induce any significant modulation of IL-1β and IL-6 expression, we observed a significant decrease in iNOS expression, which is heavily upregulated by NF-κB (Morgan and Liu, 2011). Typically, microglia cells induce iNOS expression after the exposure to inflammatory stimuli, including LPS exposure (Kim et al., 2018). The increase of iNOS expression leads to sustained production of NO and subsequent generation of reactive nitrogen oxide species (RNOS), which can mediate a broad spectrum of pathological effects, including the impairment of cell metabolism. Moreover, although NO plays several physiologic roles, including a positive modulation of synaptic transmission and brain plasticity, excessive NO generation by iNOS activation leads to the activation of inflammatory pathways and mediators, including histamine and cytokines, leading to cell damage and, ultimately, to cell death (Zamora et al., 2000). Accordingly, numerous studies document iNOS expression in a large number of human disorders (Zamora et al., 2000).

Overall, although preliminary, our data suggest that grapefruit IntegroPectin may modulate inflammatory response and basal microglia activation by inhibiting the cascade PI3K-NF-kB-iNOS.

Among the biomolecules adsorbed at the outer surface of the grapefruit pectic fibers, the flavanone glycoside naringin (4’,5,7-trihydroxyflavonone-7-rhamnoglucoside) was found to be exceptionally concentrated in grapefruit IntegroPectin (Scurria et al., 2021). Due to naringinin scavenging of free radicals activity and its effects on the inhibition of NF‐κB signaling pathway and expression of inflammatory proteins, including iNOS (Tutunchi et al., 2020), we can hypothesize a major role of this flavonone in mediating the protective effects of grapefruit IntegroPectin on microglia cells.

In conclusion, studying the protective effects of grapefruit IntegroPectin on human microglial HMC3 cells we have discovered that this new *Citrus* pectin exerts a multispectrum beneficial activity on microglia cells by inhibiting intracellular pathways typically associated with apoptotic, oxidative and neuroinflammatory response.

Coupled to successful *in vivo* tests proving the powerful cardioprotective activity of this pectin (Flori et al., 2022), these results strongly support further investigations aimed at exploring the therapeutic role of this new pectin in *in vivo* models of oxidative stress-neuroinflammatory based-diseases, including neurodegenerative disorders. Indeed, a recent study concerning the *in vivo* administration of a similar *Citrus* pectin derived via hydrodynamic cavitation from the peels of *Citrus reticulata* (mandarin) has shown immunomodulatory activity and reduction in LPS-induced lymphocytes in rats, though not linearly associated with dosage (Putri et al., 2022).

Featuring a particularly low (22%) degree of crystallinity (Piacenza et al., 2022) and a very low degree of esterification (the fraction of -COOH groups esterified with methanol) of 14% (Presentato et al., 2020), grapefruit IntegroPectin is uniquely rich in adsorbed naringin (74 mg/g) (Scurria et al., 2021) and in long and numerous hydrophilic RG-I regions (Presentato et al., 2020). All this favours the interaction of the IntegroPectin chains rich in hydrophilic carboxylate and hydroxyl groups with the HMC3 cell membrane, enhancing the delivery of bioactive molecules, including the otherwise poorly soluble naringin, well-known for its anti-inflammatory and antioxidant activity (Tutunchi et al., 2020). Taking into account the recently discovered antibacterial (Piacenza et al., 2022) (Presentato et al., 2020), cardioprotective (Flori et al., 2022), mitoprotective, antioxidant and antiproliferative properties of grapefruit IntegroPectin (Nuzzo et al., 2021b), these new findings suggest that this novel pectin derived from *Citrus paradisi* peel residues of the industrial juice extraction using water and electricity constitutes one of the single most eminent examples of pectin as universal medicine (Zaitseva et al., 2020).

## Author Information

### Co-First Authors

**Miriana Scordino** - *Dipartimento di Biomedicina, Neuroscienze e Diagnostica Avanzata, Università di Palermo, corso Tukory 129, 90134 Palermo, Italy*;

**Giulia Urone** - *Dipartimento di Biomedicina, Neuroscienze e Diagnostica Avanzata, Università di Palermo, corso Tukory 129, 90134 Palermo, Italy*

### Authors

**Giuseppa Mudò** - *Dipartimento di Biomedicina, Neuroscienze e Diagnostica Avanzata, Università di Palermo, corso Tukory 129, 90134 Palermo, Italy*;

**Monica Frinchi** - *Dipartimento di Biomedicina, Neuroscienze e Diagnostica Avanzata, Università di Palermo, corso Tukory 129, 90134 Palermo, Italy*;

**Chiara Valenza** ^-^ *Dipartimento di Biomedicina, Neuroscienze e Diagnostica Avanzata, Università di Palermo, corso Tukory 129, 90134 Palermo, Italy*;

**Angela Bonura** - *Istituto per la Ricerca e l’Innovazione Biomedica, CNR, via U. La Malfa 153, 90146 Palermo, Italy*;

**Chiara Cipollina** - *Istituto per la Farmacologia Traslazionale, CNR, via U. La Malfa 153, 90146 Palermo*;

**Rosaria Ciriminna** - *Istituto per lo Studio dei Materiali Nanostrutturati, CNR, via U. La Malfa 153, 90146 Palermo, Italy*;

**Francesco Meneguzzo** - *Istituto per la Bioeconomia, CNR, via Madonna del Piano 10, 50019 Sesto Fiorentino FI, Italy*;

**Mario Pagliaro** - *Istituto per lo Studio dei Materiali Nanostrutturati, CNR, via U. La Malfa 153, 90146 Palermo, Italy*;

## Notes

The Authors declare no competing financial interest.

## Acknowledgements

We thank Campisi Citrus (Siracusa, Italy) for the generous gift of waste grapefruit peel from which the IntegroPectin was obtained. We also thank Dr. Stefania Raimondo for the technical assistance with RT-PCR experiments. C.V. gratefully acknowledges Italy’s Ministry of University and Research and the University of Palermo for funding her PhD program (D. M. 1061/2021).

## CRediT authorship contribution

**Miriana Scordino**: Formal analysis, Investigation, Data curation, Writing - Original Draft. **Giulia Urone**: Formal analysis, Investigation, Data curation, Writing - Original Draft. **Monica Frinchi**: Formal analysis, Investigation, Data curation, Writing - Original Draft, Visualization. **Chiara Valenza**: Formal analysis, Investigation, Writing - Original Draft. **Angela Bonura**: Resources, Supervision. **Chiara Cipollina**: Resources, Writing - Review & Editing. **Rosaria Ciriminna**: Resources, Supervision. **Francesco Meneguzzo**: Resources. **Mario Pagliaro**: Resources, Writing - Review & Editing. **Giuseppa Mudò**: Resources, Supervision, Project administration, Visualization. **Valentina Di Liberto**: Conceptualization, Methodology, Validation, Formal analysis, Visualization, Writing - Original Draft, Supervision, Project administration.

